# Proteome-Wide Serology Identifies Secreted IgD against RSV Fusion Glycoprotein as a Signature of Broadly Elevated Antibody Response

**DOI:** 10.64898/2026.11.03.764056

**Authors:** Zoe Turner, Weize Tang, Yasmine Rais, Andrei P. Drabovich

## Abstract

Serological evaluation of antibody response using conventional indirect immunoassays cannot capture the full depth, diversity, and functional complexity of human polyclonal antibodies. Here, we developed mass spectrometry-based Proteome-Wide Serology (ProWiSe) assays for the identification and absolute quantification of antigen-specific human antibodies across isotypes, subclasses, and variable-region repertoires. Using ProWiSe, we measured the depth and diversity of serum antibodies to multiple antigens of respiratory syncytial virus (RSV), a pathogen associated with repeated reinfections and skewed immune response. IgG1 antibodies targeting the fusion glycoprotein F (RSVF) dominated the antibody response, with a background-corrected median level of 1,700 ng/mL across 214 healthy maternal serum samples from the general population. ProWiSe identified reproducible co-enrichment of complement component 1q complexes, revealed the usage of several dozen IGHV variable-region germline genes in the anti-RSVF polyclonal response, and suggested limited IGHV repertoire diversity in some individuals. In addition to IgG1, we discovered a complex antibody response characterized by significantly elevated levels of nearly all human immunoglobulin isotypes and subclasses, including previously unexplored IgG4 and IgD. The identification of high-affinity IgD against prefusion and postfusion RSVF, but not against other antigens studied, was the most unexpected observation, occurring in 52% of the healthy maternal serum samples. ProWiSe unambiguously identified the secreted IgD isoform P01880-1 and supported the IgD paradox of preferential λ light-chain usage in secreted IgD. Detectable anti-RSVF IgD levels (≥0.11 ng/mL) correlated with broadly elevated antibody responses across all other isotypes and subclasses, except IgE. ProWiSe assays offer a versatile platform for quantitative serological diagnostics, exploration of depth, diversity, and dynamics of polyclonal antibody response, and discovery of rare antibody isotypes.

## INTRODUCTION

Evaluation of individual and population immunity by serology testing relies on detecting specific pathogen- or antigen-specific antibodies in blood serum or biological fluids. Human serum antibodies comprise a highly diverse mixture of isotypes, subclasses, and clonotypes that collectively shape the immune response^1^. Conventional serology testing, typically based on indirect immunoassays, cannot capture the enormous heterogeneity of human polyclonal antibodies and is limited by semi-quantitative measurements, insufficient standardization, and cross-reactivity that may contribute to lower diagnostic specificity and inconsistent clinical interpretation^2, 3^. Novel assays are needed to resolve the limitations of indirect immunoassays, explore the depth, diversity, and cooperation of human antibody isotypes (IgG, IgA, IgM, IgE, IgD) and subclasses (IgG1, IgG2, IgG3, IgG4, IgA1, IgA2) in polyclonal antibody response, and investigate the dynamics of class switching during acute primary infections, chronic antigen exposure, and repeated vaccinations^4, 5^. In this study, we developed Proteome-Wide Serology (ProWiSe) assays aimed at the identification and quantification of the full depth and diversity of the human polyclonal antibody response generated against viral antigens.

Respiratory Syncytial Virus (RSV) is a common respiratory virus associated with frequent reinfections in humans^6^. RSV infections often pose serious risks for infants and account for over a quarter of infant hospitalizations, severe complications, and 100,000 deaths annually^7^. Infants initially obtain protection against RSV with maternal antibodies that last only for the first few months of life^8^. While nearly all adults have serum anti-RSV antibodies due to repeated infections, antibody levels decrease in the elderly and immunocompromised patients^9^. Measurement of the full depth and diversity of serum anti-RSV antibodies may uncover mechanisms of RSV immune evasion and offer prognostic information on RSV severity in vulnerable populations^10^.

Previous studies identified RSV fusion glycoprotein F (RSVF) as the most immunogenic protein. They also revealed high titers of anti-RSVF total IgG, IgG1, IgG2, and IgG3 after RSV infections^11^. However, the widely inconsistent levels of anti-RSVF IgG in sera (4 µg/mL^12^ to 300 µg/mL^13^) suggested potential challenges with standardization of indirect immunoassays and justified the need for alternative, more comprehensive quantitative assays.

In this study, we aimed at the development of comprehensive proteome-wide assays to enable the in-depth characterization of anti-RSV human serum polyclonal antibodies. Our specific objectives were: (i) develop untargeted and targeted ProWiSe assays against several RSV antigens; (ii) measure absolute concentrations of all isotypes and subclasses of anti-RSV polyclonal antibodies in healthy maternal serum samples from the general population; (iii) identify the proteome co-enriched with anti-RSV antibodies; (iv) evaluate the diversity of anti-RSV polyclonal antibodies through the use of immunoglobulin heavy-chain variable region (IGHV) genes; and (v) provide a comprehensive overview of the polyclonal antibody response to RSV.

## EXPERIMENTAL METHODS

### Reagents and samples

A detailed list of antigens and antibodies used in this study can be found in **Supplemental Table S1**. Briefly, recombinant RSV antigens (Sino Biological, Wayne, PA) included: (i) post-iRSVF: post-fusion glycoprotein F expressed in baculovirus-insect cells; (ii) post-hRSVF: post-fusion glycoprotein F expressed in HEK293 cells; (iii) RSVG: major surface glycoprotein G. Pre-hRSVF, a stabilized RSV prefusion glycoprotein F, was generously provided by the David Marchant Lab (University of Alberta). SARS-CoV-2 Spike protein (Sino Biological) was expressed in baculovirus-insect cells. Recombinant monoclonal antibodies included NISTmAb 8671 (NIST reference material^14^), recombinant Nirsevimab (#DVV02802) and Clesrovimab (#DVV02808; AntibodySystem, France), and Palivizumab-CP169 (Bio X Cell, Lebanon, NH). PNGaseF (#V4831) was obtained from Promega, Madison, WI. The WHO 1st International Standard for antiserum to Respiratory Syncytial Virus (#16/284) and Non WHO reference material immunoglobulin D (IgD) serum, human (#67/037) were obtained from NIBSC, Potters Bar, UK. Mouse anti-Human IgD-UNLB (#9030-01) was obtained from Southern Biotech, Birmingham, AL. Stable isotope-labelled absolute quantified peptide internal standards (SpikeTides_TQL) were synthesized, purified, and quantified by JPT Peptide Technologies, Berlin, Germany. These standards incorporated heavy-isotope-labeled lysine (¹³C₆,¹⁵N₂) or arginine (¹³C₆,¹⁵N₄) and contained a C-terminal trypsin-cleavable quantification tag (Ser-Ala-3-nitrotyrosine-Gly). Dithiothreitol, iodoacetamide, and MS-grade Trypsin (#EMS0005) were obtained from MilliporeSigma, Oakville, ON.

### Clinical specimens

The general population serum cohort comprised 214 healthy maternal serum samples. The additional 20 male serum samples were included to assess potential sex-based differences. Separate subsets of the maternal sera were used for initial characterization of the anti-RSVF antibody repertoires by untargeted MS (N=80), characterization of total serum IgD by targeted MS (N=71), and quantification of anti-RSVF human immunoglobulin isotypes and subclasses by targeted MS (N=214). Demographic characteristics of the serum samples are presented in **Supplemental Table S2**. The use of archived, deidentified serum samples previously collected from healthy individuals was approved by the University of Alberta (#Pro00104098) and the Health Research Ethics Board of Alberta (#HREBA.CC-22-0056).

### Immunoprecipitation and proteomic sample preparation

High-binding 96-well polystyrene microplates (#655074, Greiner Bio-One, Monroe, NC) were coated overnight with recombinant antigen proteins in PBS: the initial experiments with untargeted DDA MS utilized 0.5 µg antigen/well, while the subsequent experiments with the targeted SRM and untargeted DIA measurements included 0.3 µg antigen/well. The plates were washed, blocked with 5% sucrose in PBS for 2 hours and washed. Serum (20 μL) was diluted with 5% sucrose in PBS, incubated for 2 hours, and washed with 50 mM ammonium bicarbonate. Following two hours of on-plate incubation and washing, enriched antibodies were reduced with dithiothreitol, alkylated with iodoacetamide, and digested on the same microplate with trypsin (0.5 µg/well). SpikeTides_TQL peptide internal standards (500 fmol/well for the rapid targeted SRM and 300 fmol/well for other MS measurements) were spiked before digestion. Tryptic peptides were desalted and concentrated with OMIX C18 pipette tips (#A5700310K, Agilent, Santa Clara, CA).

### Direct digest of serum

Total serum proteins (10 µg) were reduced with dithiothreitol, alkylated with iodoacetamide, and digested with trypsin overnight (2 µg/sample). SpikeTides_TQL peptide internal standards (500 fmol/well) were spiked before digestion. Tryptic peptides were desalted and concentrated with OMIX C18 pipette tips.

### Untargeted DDA LC-MS

Peptides were separated using EASY-nLC II nanoLC (Thermo Scientific, Waltham, MA) with µPAC HPLC columns (50 cm, #COL-CAP050G1B; Neo110cm, #COL-NANO110NEOB; or Neo High Throughput, #COL-CAPHTNEOB, Thermo Scientific) over 5-40% acetonitrile gradient at 500 nL/min flow rate. Peptides were ionized with an EASY-Spray source and analyzed with an Orbitrap Elite mass spectrometer (Thermo Scientific). Profile MS1 scans (303-1200 m/z; 120K resolution at 400 m/z; lock mass 445.120025 m/z) were followed by top 15 Orbitrap MS/MS scans (145-1600 m/z, 15K, centroid, isolation width 2 Th) with HCD fragmentation (normalized CE 30, activation 0.1 ms, unassigned and +1 charge states rejected, min signal 5K). Other parameters included dynamic exclusion was 12 s with repeat count 1, positive mode, 2.4 kV source voltage, 290°C capillary temperature, AGC target and max ion time for FTMS (1e6 at 100 ms) and MSn (5e4 at 50 ms). Raw files were searched with MaxQuant v2.4.0. A modified FASTA database included the UniProt human canonical proteome database (20386 entries, July 7, 2022) with the additional sequences of the 44 most frequent human germline IGHV genes and alleles (>0.5% allele frequency in the population, AIRR Data Commons), 7,272 IGHV sequences of the human antibodies derived by single-cell RNA sequencing of human B cells (CoV-AbDab database), and sequences of NISTmAb, RSVF and RSVG. Modifications included constant cysteine carbamidomethylation and variable methionine oxidation, glutamine and asparagine deamidation, and peptide N-term pyroglutamic acid formation for glutamine and glutamic acid, and up to three missed cleavages for trypsin. Volcano plots were prepared with Perseus software (v2.0.11).

### Untargeted DIA LC-MS

Peptides were separated using the NanoElute2 system (Bruker, Billerica, MA) and a two-column separation method using a PepMap Neo Trap Cartridge (5 x 0.3 mm, 5 μm C18, #174500, Thermo Scientific) and PepSep C18 analytical column (100 x 0.15 mm, 1.5 μm C18, #1893483, Bruker) over 5-35% acetonitrile gradient at 450 nl/min and 40°C. Peptides were ionized with a Captive Spray 2 source and analyzed with TimsTOF Ultra2 (Bruker). Acquisition parameters included positive mode, 200 °C source temperature, 1600 V capillary voltage, 3.0 L/min dry gas, and high sensitivity detection enabled. DIA-PASEF cycles of 1.07 s included 75 ms tims accumulation and 75 ms tims ramp time, with 0.65-1.30 1/K0 ion mobility range. A single MS1 scan (100-1700 m/z) was followed by 12 MS/MS ramps (331.3-1132.3 m/z) in a total of 32 windows over ≥ +2 ion cloud (26 Da with 1 Da overlap). Bruker .d files were automatically processed with Bruker ProteoScape (BPS; v. 2026b) and dia-PASEF Spectronaut (v. 20.4), directDIA+ (Fast), 0.01 precursor and protein global q-value cutoffs, specific trypsin/P pulsar search with 2 missed cleavages, constant cysteine carbamidomethylation, and variable methionine oxidation, asparagine deamidation, protein N-term acetylation, and peptide N-term pyroglutamic acid formation for glutamine. A modified FASTA database included the UniProt human canonical proteome database (20,652 entries, January 19, 2026) with the additional 227 unique sequences of functional IGHV germline genes with their allele variants, and sequences of RSV antigens and four mAbs.

### Targeted microflow LC-SRM

Peptides were separated using ACQUITY UPLC system (Waters, Milford, MA) with a Luna Omega Polar C18 column (50 mm × 1 mm, 1.6 μm, 100 Å, #00B-4748-A0, Phenomenex, Torrance, CA) over a 5-40% acetonitrile gradient at 45 μL/min flow rate. Peptides were ionized with optiFlow Turbo V nanospray ion source and analyzed with QTRAP 6500+ (SCIEX, Marlborough, MA). Acquisition parameters included: 150°C source temperature, +4500 V electrospray ionization voltage; 35 psi air curtain gas, 35 psi for atomizing Gas 1 and 15 psi auxiliary Gas 2, 80 V declustering potential, 10 V entrance potential, 12 V collision cell exit potential, and collision gas “High”. Quantified peptides, internal standards, and SRM transitions are presented in **Supplemental Table S3.** The acquisition order of LC-MS injections (N=856) was randomized in blocks of 24 injections (12 serum samples in duplicates) of PBS negative controls followed by 2 injections of BSA, 24 injections of post-iRSVF-enriched antibodies, and 2 injections of BSA.

### Differential quantification of anti-RSV immunoglobulin isotypes and subclasses

Skyline software (v23.0.9) was used to analyze untargeted DDA MS, untargeted DIA MS, and targeted SRM data, and calculate the light-to-heavy (L/H) ratio. For DDA MS, the light endogenous and heavy isotope-labelled MS1 precursor peaks (MS1 filtering: M, M+1, M+2 precursor peaks, 120K resolving power at 400 m/z) were extracted. For dia-PASEF data, MS2 filtering included DIA, centroid, all ions, 10 ppm. For SRM data, MS2 filtering included PRM, QIT, 0.7 m/z. Serum volume (20 μL), internal standard amounts (300 or 500 fmol/well), and L/H ratios were used to calculate the absolute concentrations of each isotype and subclass of human antibodies.

### Analysis of WHO International Standard Antiserum

The WHO 1st International Standard for antiserum to Respiratory Syncytial Virus (16/284) arrived as a lyophilized powder and was re-constituted to 2000IU/mL. Onto a plate coated with post-iRSVF (300ng/well) and blocked with 5% sucrose in PBS, the antiserum was diluted 1/5 (20μl per well), incubated for 2 hours and washed with 50 mM ammonium bicarbonate. Enriched antibodies were reduced with dithiothreitol, alkylated with iodoacetamide, and digested on the same plate with trypsin (0.5 µg/well). SpikeTides_TQL peptide internal standards (300 fmol/well) were spiked before digestion, and samples were run using the untargeted DIA LC-MS methods. The absolute concentration of each isotype and subclass was determined; the dominant isotypes/subclasses (IgG1, IgG2, IgA1, IgA2, and IgM) concentrations were summed and divided by 2000 IU/mL to get an approximate mass of antibody per 1 IU.

### Sample size and power calculations

Power calculations were performed using G*Power (v3.1.9.6; Universität Düsseldorf) for two-tailed Wilcoxon signed-rank tests of matched pairs (α=0.05). For 214 paired samples analyzed by microflow LC-targeted SRM, the post hoc achieved power was >99.9% for IgG1 and 92% for IgD, with effect sizes of 1.08 and 0.23, respectively. Effect sizes were calculated as the mean paired post-iRSVF versus PBS differences divided by the standard deviation of the paired differences (1,900±1,765 ng/mL for IgG1 and 8.40±36.3 ng/mL for IgD). To achieve 80% power, the required sample sizes were 10 matched pairs for IgG1 and 157 for IgD.

## RESULTS

### ProWiSe assay development

ProWiSe assays can be considered as indirect immunoassays in which affinity-enriched human antibodies are analyzed with mass spectrometry (MS) rather than with a secondary anti-human antibody (**Fig. 1**). Each isotype (IgG, IgA, IgM, IgE, IgD) and subclass (IgG1, IgG2, IgG3, IgG4, IgA1, IgA2) of human immunoglobulins has unique trypsin-cleaved peptides within the heavy constant regions, and such peptides can be utilized for unambiguous identification and differential quantification. ProWiSe assays rely on the specific and high-affinity recognition of an antigen by serum polyclonal antibodies, with the identical setup of affinity enrichments for indirect immunoassays and ProWiSe assays. Affinity enrichment and proteomic sample preparation (denaturation and digestion of antibodies, antigens, and associated proteins) are performed in the same 96-well microplate to simplify sample processing, minimize protein loss, facilitate automation, increase throughput, and improve reproducibility. ProWiSe assays can be employed for either unbiased discovery and characterization of human polyclonal antibodies using extended nanoflow liquid chromatography separations and deep untargeted MS analysis, or absolute quantification and validation of the complete panel of human immunoglobulin isotypes and subclasses in large patient cohorts using rapid microflow separations, targeted MS analysis, and internal standards^3^.

**Figure 1.**
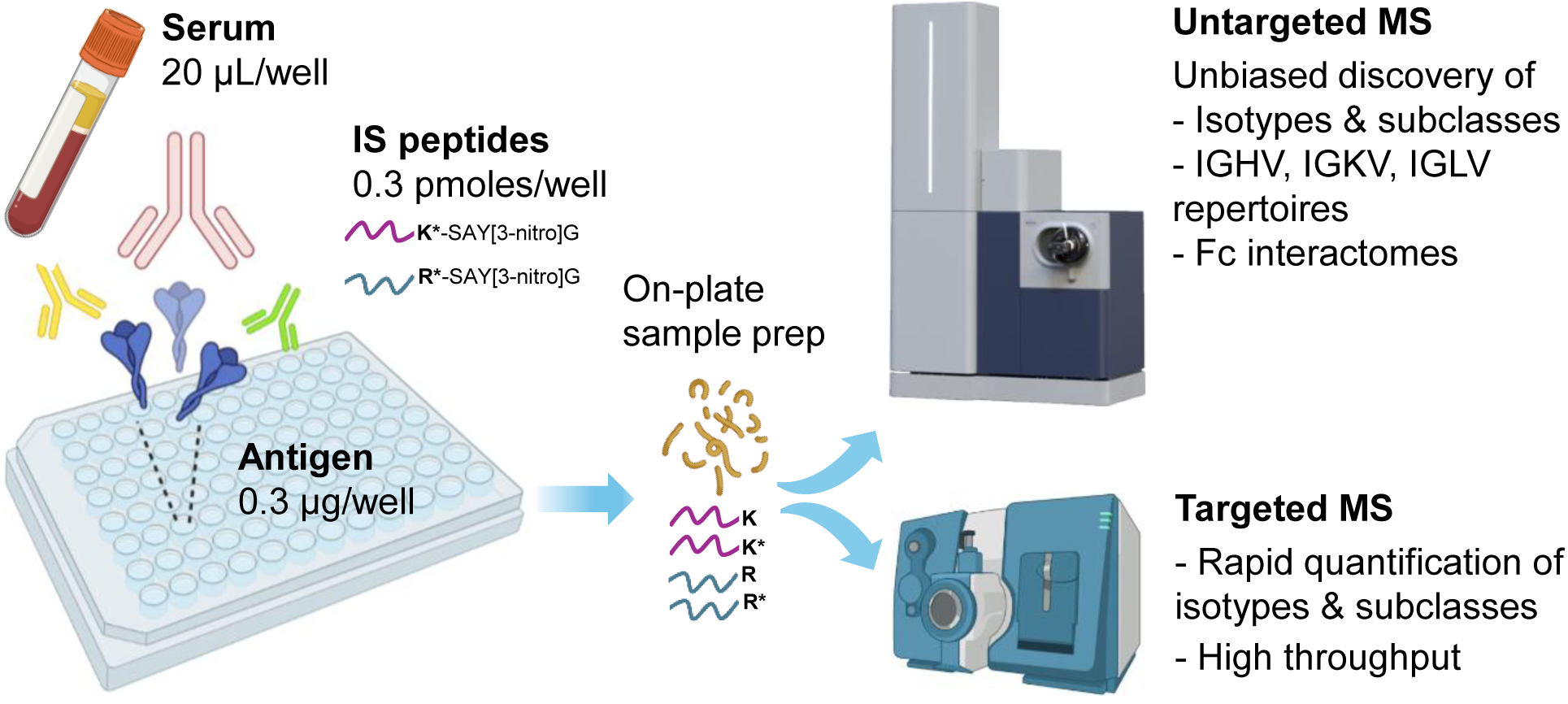
The setup of untargeted and targeted ProWiSe assays.

In this study, the sixteen carefully designed, synthesized, and purified peptide internal standards facilitated absolute quantification with high analytical selectivity and reproducibility (**Supplemental Table S4**). These peptides incorporated heavy-isotope-labeled lysine or arginine and contained a C-terminal trypsin-cleavable tag to standardize tryptic digestion, along with 3-nitrotyrosine for absolute quantification by spectrophotometry^15^. The heavy-isotope labels allow for calculation of the light-over-heavy (L/H) ratio and quantification at MS1 precursor level with untargeted data-dependent acquisition (DDA) workflows, and at MS2 fragment level with untargeted data-independent acquisition (DIA) and targeted selected reaction monitoring (SRM) workflows. To design and develop these peptides, we used our in-house proteomics data^3^ and public NextProt database^16^ to select the most abundant, unique, and fully tryptic peptides of the heavy chain constant regions for each isotype and subclass of human immunoglobulins. Peptides with known post-translational modifications (PTMs) were excluded. Data mining of a population-wide genomic database (https://gnomad.broadinstitute.org; ∼388,000 individuals and 776,000 alleles) excluded peptides with high-frequency missense variants and allotypes. For example, the most frequent variant for the IGHG1 tryptic peptide K.GPSVFPL**<u>A</u>**PSSK.S, p.Ala12Val, was present with allele frequency of only 1.3e-5 (**Supplemental Table S4**). Finally, linear correlation of MS1 area of several peptides per protein in 10 patient sera was used to exclude peptides with poor performance due to potentially poor stability, in-source fragmentation, LC separation issues, and unknown PTMs and variants. The peptide purity and efficiency of trypsin digestion were also measured (**Supplemental Table S5**). An absolute average amount of post-iRSVF antigen coated onto a microplate well was 91±18 ng, as measured by MS. Thus, ProWiSe assays reported the absolute concentration of human antibody isotypes and subclasses enriched from serum and ensure assay standardization across hundreds of patient serum samples.

### Evaluation of ProWiSe assays with anti-RSVF monoclonal antibodies

To validate ProWiSe assays, we utilized a high-affinity interaction between recombinant post-iRSVF protein and monoclonal antibody NISTmAb (K_d_ = 2.7 nM^17^). NISTmAb, a certified NIST reference material 8671^14^, is a recombinant humanized IgG1 kappa immunoglobulin composed of two identical light and heavy chains, with heterogeneous glycosylation forms, N-terminal pyroglutamination, and C-terminal lysine clipping^18^. We demonstrated that the simple affinity capture of NISTmAb on microplates resulted in 640-fold enrichment compared to a no-antigen control **(Fig. 2A-D)**. Additional recombinant monoclonal antibodies, palivizumab (post-fusion K_d_=1.4 nM^19^), clesrovimab (post-fusion K_d_=0.48 nM^20^), as well as the pre-fusion-specific nirsevimab ( K_d_=0.12 nM^21^), were used as positive controls for ProWiSe assays and validation of pre-fusion and post-fusion forms of the RSVF antigen **(Figure 4A)**.

**Figure 2.**
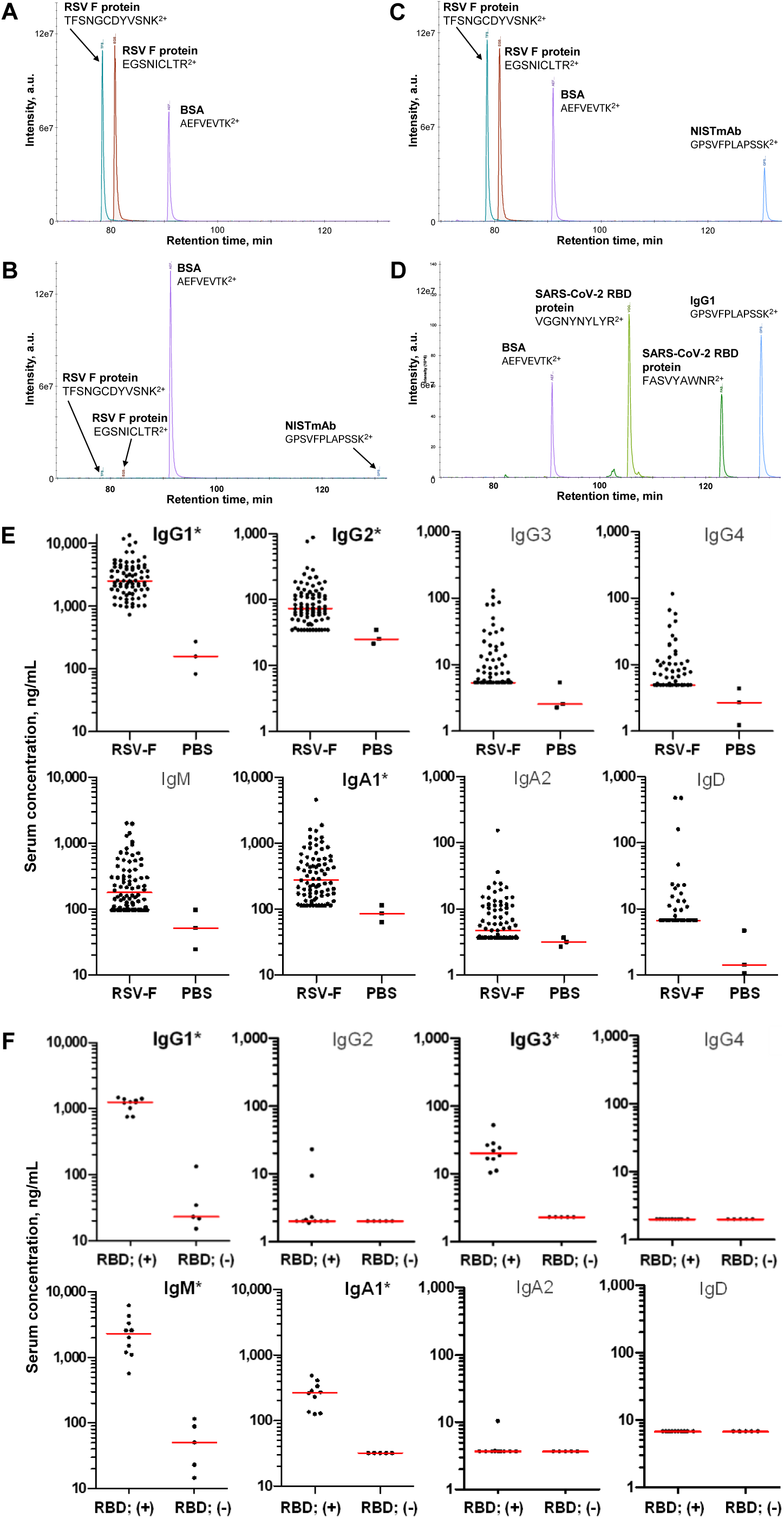
Enrichment of NISTmAb 8671 and endogenous polyclonal antibodies with an RSVF protein, as measured by ProWiSe assay with label-free MS1 quantification. **(A)** Positive control demonstrating that post-iRSVF protein (0.5 µg/well) is coated onto the high-binding immunoassay microplates. Two tryptic RSVF peptides were measured, with a BSA peptide was presented as a loading control. **(B)** Negative control demonstrating the lack of non-specific binding of NISTmAb 8671 monoclonal antibody (1 µg/well) to the microplates coated with PBS and blocked with BSA **(C)** ProWiSe assays revealed that NISTmAb (1 µg/well) was enriched 640-fold with post-iRSVF protein (0.5 µg/well) relative to the negative control. **(D)** As an additional positive control, SARS-CoV-2 RBD protein (0.5 µg/well) enriched IgG1 antibodies from a pool of convalescent human sera (20 µL/well), as previously reported^3^. NISTmAb, an IgG1 antibody, shares a unique tryptic peptide, GPSVFPLAPSSK, with endogenous human IgG1. **(E)** RSV-specific polyclonal antibodies were enriched from 80 maternal sera samples using an RSV post-iRSVF antigen, with three matched no-antigen negative controls. **(F)** For comparison, SARS-CoV-2-specific polyclonal antibodies were enriched with an RBD antigen from 10 PCR-confirmed convalescent plasma samples (+) and 5 pre-pandemic negative plasma (-). All IA enrichments and trypsin digestions (A-F) were completed onto a single microplate, and nanoLC-MS/MS with label-free MS1 quantification were used for analysis. *indicated statistically significant isotypes or subclasses determined using Bonferroni-adjusted 1-tail Mann-Whitney U test *P*-values < 0.0063. LOD levels were calculated in positive samples based on detectable peaks of the light endogenous peptides with a signal-to-noise ratio>3 for the combined MS1 intensity of a precursor and M+1 forms. Concentrations below LOD were adjusted to the LOD levels. PBS group levels below LOD are presented to demonstrate the variability of non-specific binding for all isotypes. PBS, no-antigen negative control with PBS buffer; RSVF, postfusion RSVF expressed in baculovirus-insect cells; RBD (+), Receptor binding domain of the spike glycoprotein SARS-CoV-2; RBD(-), no-antigen negative control with PBS buffer.

### Identification and quantification of isotypes and subclasses of human anti-RSV antibodies

We previously demonstrated the use of highly sensitive and selective immunoaffinity-targeted MS assays as an experimental approach for the rational design of serological diagnostics^3^. We measured relatively high absolute concentrations (1-2 µg/mL IgG1) of viral antigen-specific antibodies in convalescent sera^3^. These relatively high anti-viral antibody levels motivated us to extend our assays to less sensitive but more comprehensive untargeted proteomic analyses to enable unbiased discovery, characterization, and quantification of human polyclonal antibodies. To characterize the post-iRSVF antibody response using untargeted proteomics, we analyzed an initial cohort of 80 healthy maternal serum samples from the general population. In this study, a median IgG1 concentration of 2,530 ng/mL against post-iRSVF antigen was determined **(Fig. 2E, Supplemental Table S6)**. As a positive control, antibodies against SARS-CoV-2 RBD were measured in SARS-CoV-2 convalescent sera and revealed a median IgG1 concentration of 1,250 ng/mL **(Fig. 2F, Supplemental Table S7)**. Low levels of anti-RSVF IgM (median 177 ng/mL) revealed the absence of recent RSV infections, in contrast to high levels of anti-SARS-CoV-2 RBD IgM (median 2,300 ng/mL) in SARS-CoV-2 convalescent sera. Antigen-specific serum IgE has never been detected with either RSV or SARS-CoV-2 antigens in our studies. Apart from IgE, anti-RSVF antibodies revealed elevated levels of all other immunoglobulin isotypes. This included antigen-specific IgG4 and IgD, which were not previously observed in acute SARS-CoV-2 infections^3^. Such diversity of isotypes and subclasses motivated us to further investigate the polyclonal antibody response to RSV antigens.

### Co-enrichment of Fc interactome

Unbiased ProWiSe assay revealed the Fc interactome, such as proteins associated directly and indirectly with the enriched anti-RSVF antibodies (**Fig. 3A**). Reproducible enrichment of the complement component 1q complex (C1qA, C1qB, and C1qC protein subunits) and complement cascade components (C4B) were identified. The co-enrichment of C1q and C4B was consistent with complement-dependent cytotoxicity of anti-RSVF IgG1 and IgG3^22^. Likewise, reproducible enrichment of CD5L and a linear correlation between CD5L and IgM levels supported the presence of covalent CD5L-IgM complexes, as recently reported (**Fig. 3B**)^23, 24^. It should also be noted that the soluble forms of high-affinity Fc receptors were not identified.

**Figure 3.**
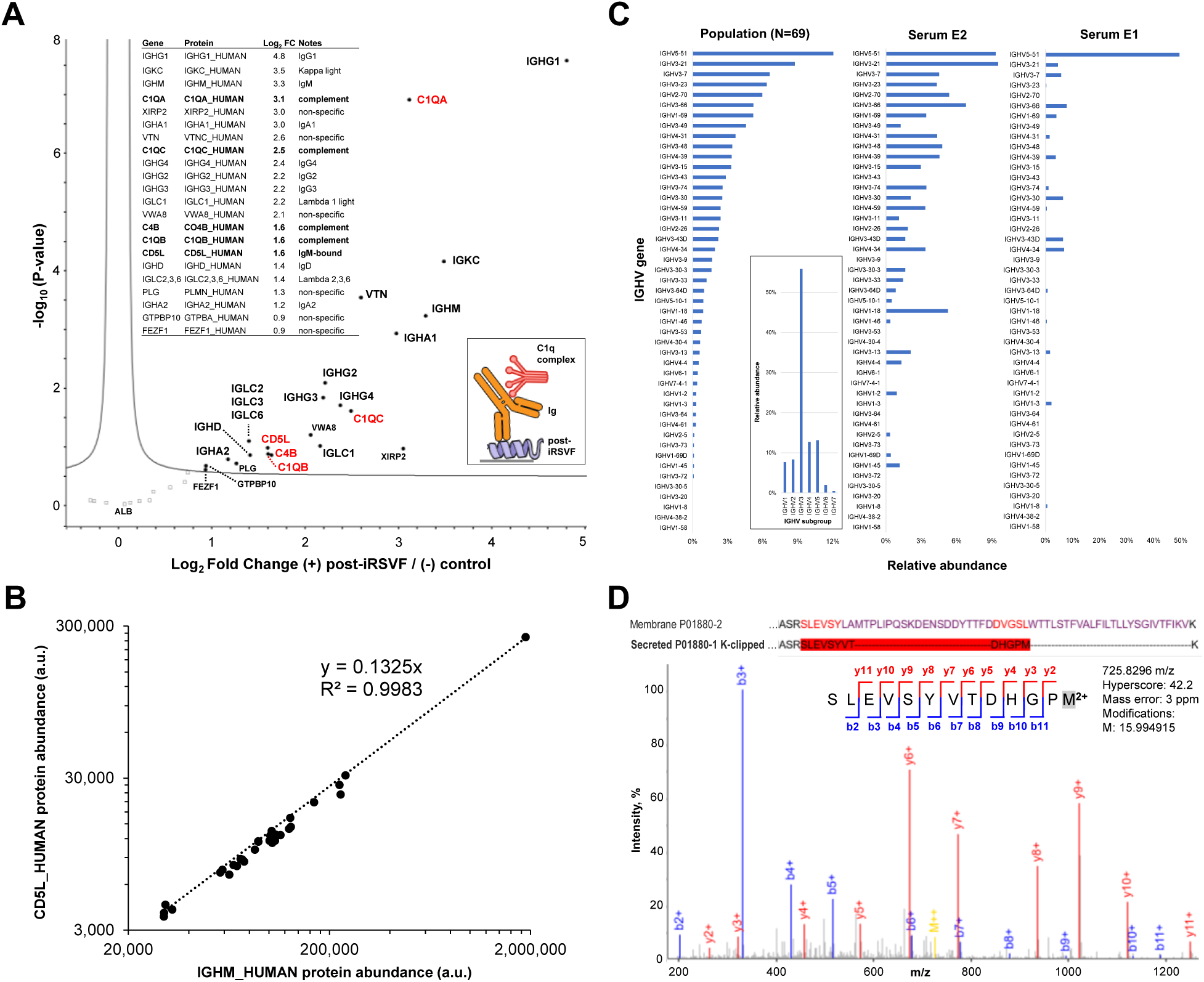
Characterization of anti-RSVF human antibodies with ProWiSe assays. **(A)** Identification of the interactome co-enriched with anti-RSVF human antibodies. Untargeted ProWiSe analysis identified human immunoglobulin isotypes and subclasses, kappa and lambda light-chain constant regions, and immunoglobulin-associated proteins including the C1q complement complex subunits C1QA, C1QB, and C1QC, C4B of the classical complement pathway, and CD5L known to covalently bind IgM. Polyclonal antibodies were enriched from 62 serum samples using post-iRSVF protein and from three matched serum samples without an antigen (negative control with PBS buffer only). The insert presents gene and protein names, and corresponding log2-transformed fold changes (log2 FC) estimated from MaxQuant LFQ values. **(B)** The reproducible enrichment of CD5L and strong linear correlation of its abundance with IgM levels suggested the presence of covalent CD5L-IgM complexes. **(C)** IGHV repertoire profiling of polyclonal anti-post-iRSVF antibodies. MaxQuant iBAQ values for each identified IGHV gene were normalized to IgG1 iBAQ values, and relative abundance was calculated as the sum of normalized iBAQ values for each subgroup divided by the total normalized iBAQ signal. The relative abundance of IGHV genes across the study population revealed sera with the highest (E2) and the lowest (E1) repertoire diversity. Although 23 genes contributed to the most abundant subgroup IGHV3 (56% of total iBAQ intensity), the most abundant single gene was IGHV5-51. **(D)** Identification of secreted C-term/lysine-clipped isoform of IgD. Untargeted ProWiSe assay identified a lysine-clipped peptide SLEVSYVTDHGPM unique for a secreted isoform P01880-1 of IgD. Peptide identification was performed using a Bruker TimsTOF Ultra 2 and MSFragger.

### IGHV repertoire profiling of anti-RSVF polyclonal antibodies

Recent studies have revealed that the polyclonal antibody response was dominated by only a small number of immunoglobulin heavy-chain variable (IGHV), immunoglobulin kappa light-chain variable (IGKV), and immunoglobulin lambda light-chain variable (IGLV) germline genes^25^. Here, we focused on IGHV germline genes that encoded the major structural scaffolds of polyclonal antibodies and investigated whether ProWiSe assays could identify the diversity and depth of IGHV repertoires in the polyclonal antibody response. To identify the IGHV genes within a patient’s antibody repertoire, we generated an in-house database that included the UniProt human proteome, 44 sequences of the most frequent germline IGHV alleles (>0.5% frequency in the population based on AIRR-seq data^26^), and 7,272 IGHV sequences of the anti-SARS-CoV-2 human antibodies (CoV-AbDab single-cell RNA-seq data^27^). The use of anti-SARS-CoV-2 sequences to search anti-RSV data facilitated the identification of IGHV alleles through shared unmutated IGHV sequences and some framework region peptides with shared point mutations, while the hypervariable CDR regions were excluded from the search.

To discover the IGHV repertoires, anti-RSVF polyclonal antibodies were enriched from maternal serum samples and analyzed by ProWiSe. A total of 2,771 MS/MS spectra, 556 unique IGHV peptides, and ∼400 distinct IGHV sequences representing all IGHV subgroups were identified in 69 serum samples (**Fig. 3C**). The IGHV3 subgroup was the most abundant based on MaxQuant intensity-based absolute quantification (iBAQ). Overall, we identified 47 unique IGHV germline genes, with 50% of the total iBAQ intensity represented by 7 unique genes. The sample with the highest diversity of IGHV repertoires had 31 unique IGHV genes identified, with 50% represented by 8 unique genes. The sample with the lowest repertoire diversity had 18 unique IGHV genes identified, with 50% represented by a single germline gene IGHV5-51. ProWiSe assays may thus enable screening of patient sera for high-affinity low-diversity oligoclonal antibodies and pave the way for *de novo* sequencing of human antibodies enriched directly from serum.

### Serum anti-RSVF IgD is present exclusively as the secreted isoform

The identification of secreted high-affinity serum IgD (immunoglobulin D) against post-iRSVF was the most unexpected discovery (**Fig. 2E**). Discovered in the 1960s, IgD remains the most mysterious immunoglobulin isotype^28, 29^. IgD is one of the least abundant immunoglobulins, with a median level of 30 µg/mL in adult serum^30^. The human IGHD gene encodes the constant heavy chain of IgD and has two alternatively spliced isoforms. A membrane-bound IGHD_HUMAN protein (P01880-2; 430 aa) is an integral part of a B cell antigen receptor, while P01880-1 (384 aa) corresponds to a secreted isoform. Previous immunoassay-based studies could not distinguish between these two isoforms and were describing the ‘soluble’ serum IgD. Here, ProWiSe assays unambiguously identified a C-terminal and lysine-clipped peptide SLEVSYVTDHGPM, unique to a secreted isoform (**Fig. 3D**). Conversely, a peptide unique to the transmembrane isoform, P01880-2, SLEVSYLAMTPLIPQSK, previously identified in B cell proteomics studies^31^, has not been identified in this study. Thus, our findings indicated that anti-post-iRSVF serum IgD was present exclusively as the secreted IgD isoform P01880-1. In the general population sera, the secreted anti-post-iRSVF IgD was identified in both maternal sera (**Fig. 2E**) and male sera (**Supplemental Fig. S1**). Although secreted anti-RSVF IgD has not, to our knowledge, been previously identified in immune cell models or in human serum, secretion of polyreactive anti-RSVF IgM by regulatory and plasma B cells has previously been reported^10^. This finding motivated us to evaluate other RSV antigens for their IgD enrichment.

### Secreted IgD recognizes RSVF but not surface glycoprotein G

We previously evaluated the responses of human antibody isotypes and subclasses to four SARS-CoV-2 antigens (S, S1, RBD, and N) in convalescent sera of patients who had recovered from SARS-CoV-2^3^. No SARS-CoV-2-specific IgD was detected in our previous studies or with the improved ProWiSe assays (**Fig. 2F)**. Here, we first measured the isotype and subclass specificity of serum polyclonal antibodies against post-iRSVF and the RSV surface glycoprotein G (RSVG). RSVG antigen substantially enriched IgG1, IgG2, and some other isotypes but not IgD (**Supplemental Table S8)**.

### Polyclonal secreted IgD recognizes both pre-fusion and post-fusion forms of RSVF

Endogenous RSVF protein is known to exist in two conformations, a metastable prefusion form and a stable postfusion form^32, 33^. Recent advances in RSV prevention have been attributed to the development of neutralizing monoclonal antibodies against the prefusion form, such as nirsevimab targeting the prefusion-specific antigenic site Ø^6, 34^. Our preliminary studies were completed with post-iRSVF generated with the baculovirus-insect expression system. Initial tests with a commercially available prefusion RSVF revealed poor binding of nirsevimab (K_d_ >1200 nM, in agreement with previous studies^33, 35^) and suggested a predominantly postfusion RSVF conformation. The stabilized prefusion RSVF form L305I DS-Cav1^36^ (referred to as pre-hRSVF) enriched nirsevimab, and the presence of prefusion and postfusion RSVF forms was confirmed by binding to a panel of monoclonal antibodies (**Fig. 4A)**. Finally, we simultaneously enriched and measured the polyclonal antibodies against several forms of RSVF including the postfusion forms from baculovirus-insect (post-iRSVF), HEK-293 (post-hRSVF) and pre-fusion stabilized RSVF (pre-hRSVF). We also evaluated RSVG and SARS-CoV-2 spike glycoprotein S (**Fig. 4B**). The secreted high-affinity IgD was reproducibly enriched with all forms of the RSVF antigen, but not with RSVG or SARS-CoV-2 S antigens, thus indicating specific affinity of serum IgD to RSVF.

**Figure 4.**
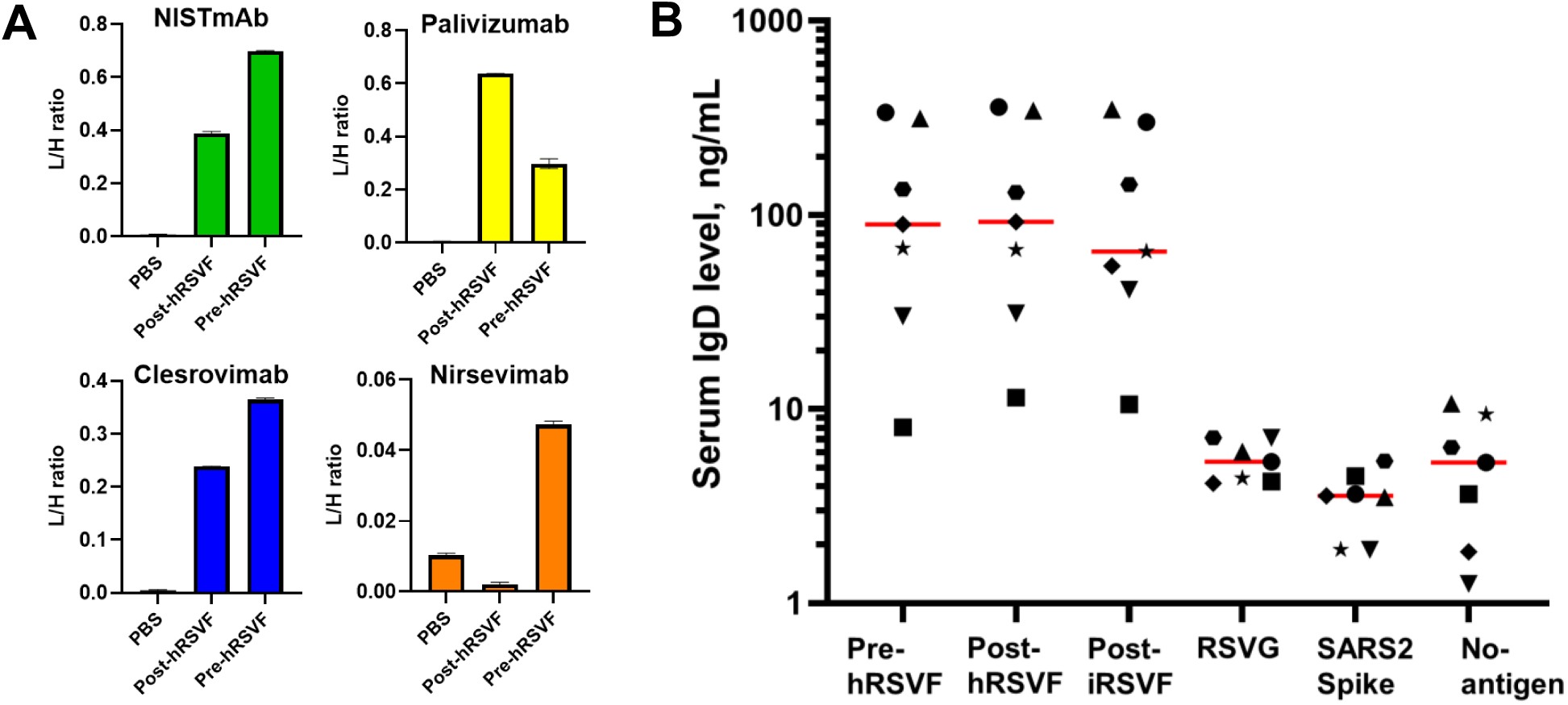
Serum IgD binds to the prefusion and postfusion RSVF. **(A)** The presence of a stabilized prefusion form was confirmed with binding to a panel of monoclonal antibodies. Nirsevimab, which recognizes antigenic site Ø that is present exclusively in the prefusion conformation, bound the pre-hRSVF antigen. **(B)** Serum IgD specifically bound to prefusion and postfusion F expressed in human HEK293 cells (pre-hRSVF and post-hRSVF), and to postfusion F expressed in baculovirus-insect cells (post-iRSVF). No binding was found for RSVG antigen expressed in HEK293 or SARS-CoV-2 Spike expressed in baculovirus-insect cells, relative to the no-antigen PBS negative control. Individual serum samples are indicated by distinct symbols for each of the tested antigen groups. A Friedman nonparametric repeated-measures test demonstrated an overall difference among groups (*P=*0.00003), followed by Dunn’s multiple-comparisons test. IgD binding to pre-hRSVF (adjusted *P=*0.02), post-hRSVF (adjusted *P=*0.003), and post-iRSVF (adjusted *P=*0.008) was significantly higher than the no-antigen control.

### Lambda light chains dominate in secreted IgD

It was previously reported that myeloma IgD and serum IgD from healthy individuals were preferentially associated with λ light chains (∼60–90% of IgD), whereas transmembrane IgD predominantly contained κ light chains, a phenomenon known as the ‘IgD paradox’^28^. Here, we used a mouse IgG2aκ monoclonal antibody against human IgD to enrich secreted IgD from three human serum samples, and analyzed IgD light-chain composition. We ensured that only tryptic peptides unique to human IGKC_HUMAN and not shared with IGKC_MOUSE were used for quantification. As a result, human IgD was enriched 440-fold. Although low levels of other human immunoglobulins were detected (IgG1 was ∼20-fold less abundant than IgD), their contribution to the κ/λ ratio was insignificant. The enriched IgD had an average κ/λ ratio of 0.59, while the average κ/λ ratio in anti-post-iRSVF polyclonal antibodies was of 1.78 (N=69; **Supplemental Figure S2**). Our measurements demonstrated that ∼63% of light chains associated with secreted IgD were λ light chains, consistent with the ‘IgD paradox’. These findings warrant further investigation into whether secreted λ-enriched IgD and transmembrane κ-enriched IgD may represent distinct forms of IgD with potentially different functional roles.

### Reference intervals of anti-RSVF polyclonal antibodies in human serum

To validate anti-RSVF antibody levels and isotype diversity in a larger population and estimate the prevalence of anti-RSVF IgD, we developed a rapid multiplex selected reaction monitoring (SRM) assay. Simultaneous 96-well plate immunoaffinity enrichments coupled to online microextraction and microLC separations enabled measurement of anti-post-iRSVF antibodies at a throughput of 120 runs/day (**Fig. 5A**). This assay enabled reproducible quantification of anti-RSVF antibodies in 214 healthy maternal serum samples, together with matched no-antigen negative controls, across 856 LC-MS injections. (**Fig. 5B, Supplemental Table S9**). The levels of immunoglobulin isotypes and subclasses were significantly elevated in the RSVF group as compared to the no-antigen negative control group (**Table 1**). To calculate reference intervals^37^ for anti-RSVF isotypes and subclasses in serum (**Supplemental Table S10**), the non-specific binding levels measured in the PBS controls were subtracted (**Supplemental Table S11**). As a result, anti-RSVF IgD was detectable in 52% of healthy maternal serum samples.

**Figure 5.**
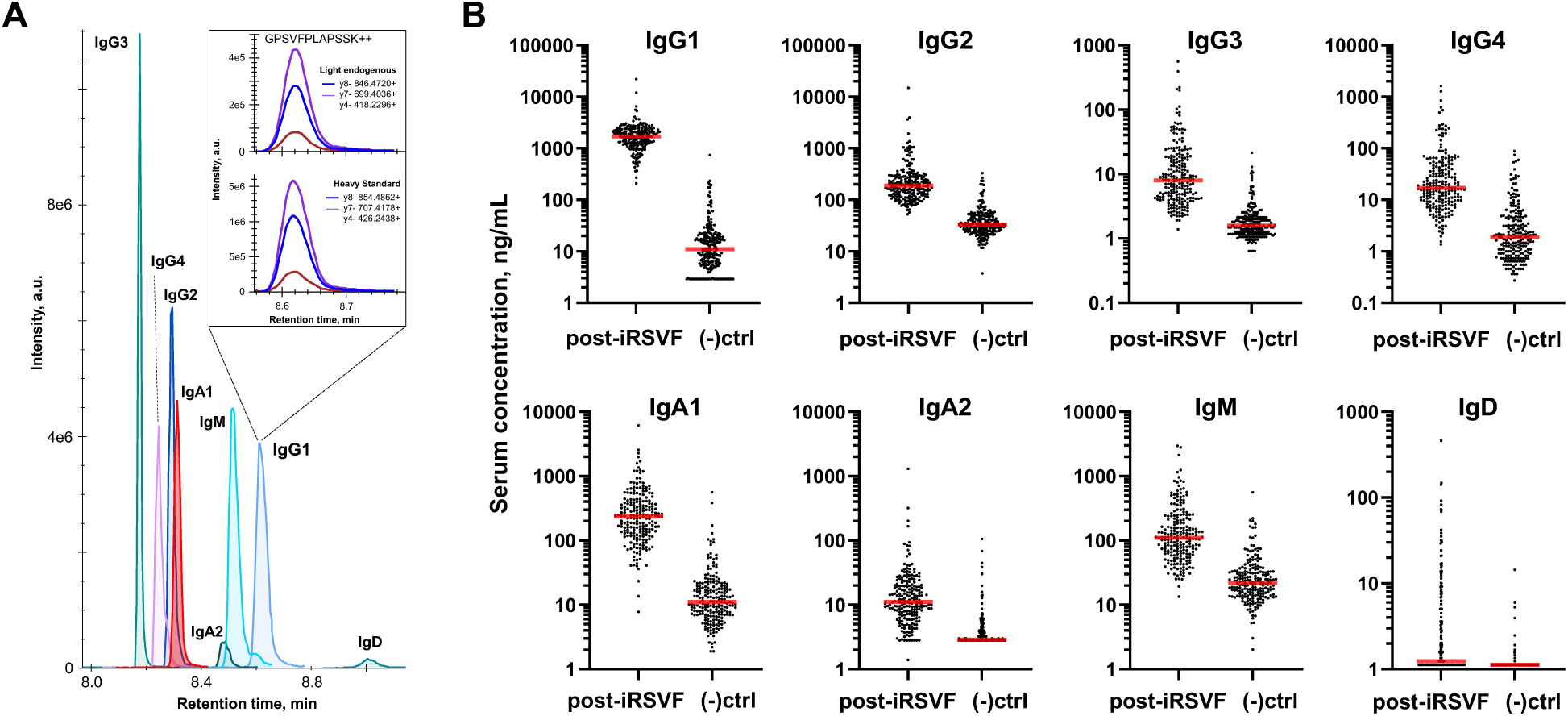
Identification, quantification, and characterization of the endogenous polyclonal anti-post-iRSVF antibodies by rapid targeted LC-SRM assay. **(A)** The rapid LC-SRM assay provided multiplexed quantification of human antibody isotypes and subclasses with high throughput (>120 runs per day). Heavy isotope-labeled peptide standards with trypsin-cleavable tags enabled ‘absolute quantification’ based on the light-to-heavy peptide ratios. **(B)** Polyclonal antibodies were enriched from 214 maternal sera samples with either post-iRSVF antigen or no-antigen controls (-ctrl). The LOD for each isotype and antibody was determined as the lowest L/H ratio producing a detectable light peptide peak with a signal-to-noise ratio >3. Concentrations below LOD were adjusted to the LOD levels. The LOD of IgD was 1.13 ng/mL.

**Table 1.** Serum concentrations of anti-RSVF immunoglobulin isotypes and subclasses measured by IA-targeted SRM. A cohort of 214 healthy maternal serum samples from the general population was analyzed using either a post-iRSVF antigen (+RSVF) or no-antigen negative control (–PBS). To calculate statistical significance, concentrations below LOD were adjusted to the LOD levels, with *P* = 0.05 used as the significance threshold for a 2-tailed Wilcoxon signed-rank (WSR) test for matched samples was used as a statistically significant cut-off. IgE isotype was not detected in any samples. Median concentrations and interquartile ranges [IQR] are presented. Reference intervals were calculated from background-corrected concentrations following subtraction of matched PBS values.

|  | **IgG1** | **IgG2** | **IgG3** | **IgG4** | **IgM** | **IgA1** | **IgA2** | **IgD** |
|---|---|---|---|---|---|---|---|---|
| (+) RSVF, ng/mL | 1,701 | 188 | 7.9 | 17 | 111 | 236 | 11 | 1.2 |
| [IQR] | [1,242-2257] | [123-282] | [4.3-18] | [9.4-41] | [64-217] | [130-398] | [7.1-20] | [1.1-5.0] |
| (–) PBS, ng/mL | 11 | 33 | 1.6 | 1.9 | 22 | 11 | 2.9 | 1.1 |
| Fold Change | 154 | 5.7 | 5.0 | 8.9 | 5.1 | 21 | 3.8 | 1.1 |
| WSR *P*-value | 7.3E-37 | 5.7E-36 | 5.4E-34 | 7.4E-32 | 2.4E-34 | 9.4E-36 | 1.0E-27 | 3.3E-18 |
| Background-corrected reference intervals in healthy maternal sera, ng/mL | 430 - 4,832 | 0 - 1,285 | 0 - 163 | 0 - 506 | 1.0-1,045 | 7.9 - 1,647 | 0 - 87 | 0 - 78 |

To facilitate inter-laboratory comparison of anti-RSVF antibody levels, we measured anti-RSVF antibodies in the 1st International Standard 16/284 for antiserum to Respiratory Syncytial Virus^38^. The standard was prepared from a pool of serum samples of six healthy adults with natural RSV infections and contained 2,000 IU/mL of neutralizing activity against RSV subtype A. Our measurements identified IgG1 as the most abundant anti-post-iRSVF isotype, at a concentration of 5,885±180 ng/mL (**Supplemental Figure S3**). Based on these measurements, 1 IU of neutralizing activity corresponded to 3.82 ng of the combined IgG1, IgG2, IgA1, IgA2, and IgM antibodies.

### Anti-RSVF IgD shows limited correlation with the total serum IgD

Previous measurements reported low levels of total soluble IgD in human adult serum (median 30 µg/mL)^39^ compared to other antibody isotypes, and wide normal ranges in adults varying from 2-37 µg/mL^40^ to 7-170 µg/mL^39^, 10-112 µg/mL^41^, and 5-240 µg/mL^42^. In IgD monoclonal gammopathy of undetermined significance or multiple myeloma, serum IgD was detected at levels exceeding 2 mg/mL^43^. Here, we measured the total serum IgD and anti-post-iRSVF IgD in 71 matched sera with previously detectable anti-post-iRSVF IgD (**Fig. 6A**). Total serum IgD (median 87 μg/mL) was found to be ∼4 orders of magnitude higher than RSVF-specific IgD (median 9.3 ng/mL). Thus, anti-RSVF IgD represented only a small fraction of total serum IgD, with limited correlation between their levels.

**Figure 6.**
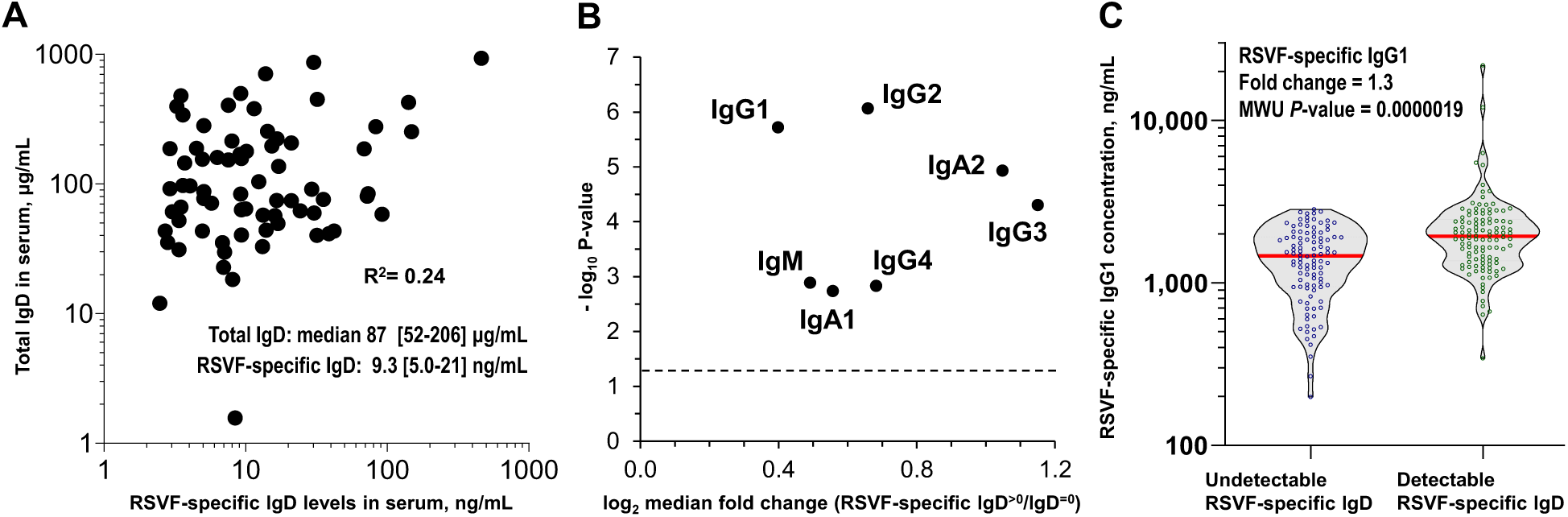
Correlations of serum RSVF-specific IgD. **(A)** Analysis of 71 matched serum samples revealed limited correlation between the total serum IgD and post-iRSVF-specific IgD. The total serum IgD concentration was found ∼4 orders of magnitude higher than RSVF-specific IgD concentrations. **(B)** Levels of human isotypes and subclasses against anti-post-iRSVF were compared between maternal serum samples with detectable RSVF-specific IgD (background-corrected IgD≥0.11 ng/mL; N=112) and samples with undetectable IgD (background-corrected IgD<0.11ng/mL; N=102). The samples with detectable RSVF-specific IgD were associated with statistically significant higher levels of all measured antibody isotypes and subclasses (Mann-Whitney U test, *P*-value < 0.05). **(C)** Violin plot showing distribution of IgG1 in serum samples with detectable RSVF-specific IgD (background-corrected IgD≥0.11 ng/mL; N=112) and undetectable RSVF-specific IgD (background-corrected IgD<0.11; N=102). MWU, Mann-Whitney U test.

In addition, we measured the total IgD in Non-WHO Reference Material Immunoglobulin D (IgD) Serum, human (67/037)^44^. This reference material was prepared from plasma with high IgD levels and included 1.41 µg of IgD per unit of IgD activity^44^. Taking into account plasma and serum dilutions^45^, IgD concentration in the original 67/037 pool was estimated to be 180 µg/mL (see **Supplemental Table S12** for calculations). Surprisingly, our untargeted MS analysis of this Reference Material did not detect any unique IgD peptides. However, numerous unique peptides of IgG1, IgG2, IgG3, IgG4, IgM and IgA1 were detected. These results highlighted the need for improved IgD reference materials and their independent validation with MS-based assays.

### Detectable IgD is correlated with broadly elevated levels of anti-RSVF antibody isotypes and subclasses

Elevated production of antigen-specific IgG and IgM antibodies was previously observed in mice bearing IgD-producing plasmacytomas, suggesting a potential association between IgD production and enhanced antibody response^46^. In addition, binding of secreted antigen-specific IgD to basophils through the CD44-binding protein galectin-9 enhanced IL-4 production and augmented the Th2-type humoral response, resulting in elevated mouse IgG1 levels (a functional equivalent of human IgG4^47^). Given that several of IgD immune functions, such as binding to basophils, were evolutionarily conserved, IgD was suggested to be a part of an ancestral surveillance system that promoted immune activation and enhanced local and systemic immune responses^48^.

Here, we investigated correlations of detectable IgD and the levels of anti-post-iRSVF antibody isotypes and subclasses (**Supplemental Table S11**). Comparison of antibody concentrations between populations with detectable IgD (background-corrected IgD ≥0.11 ng/mL; N=112) and undetectable IgD (background-corrected IgD <0.11 ng/mL; N=102) revealed that detectable IgD was associated with broadly elevated antibody levels across all isotypes and subclasses except IgE (**Table 2** and **Figs. 6B, C**). It should be emphasized that a block randomization strategy was applied during sample analysis, and that IgG1 and IgG3 levels remained significantly higher (MWU *P*= 0.0461 and 0.0290, respectively) in patients with detectable IgD after adjustments for MS acquisition order. Elevated levels of RSVF-specific IgG4 could indicate amplification of Th2-associated humoral response, as previously observed in mice^47^.

**Table 2.** Levels of anti-post-iRSVF human immunoglobulin isotypes and subclasses in populations with undetectable and detectable IgD. Detectable IgD (background-corrected IgD≥0.11 ng/mL) was associated with broadly elevated antibody responses across all isotypes and subclasses, except IgE. MWU, 2-tailed Mann-Whitney U test, with *P* = 0.05 used as the significance threshold.

|  | **IgG1** | **IgG2** | **IgG3** | **IgG4** | **IgM** | **IgA1** | **IgA2** |
|---|---|---|---|---|---|---|---|
| Undetectable IgD (N=102) | 1,469 | 117 | 3.3 | 12 | 66 | 160 | 5.2 |
| Detectable IgD (N=112) | 1,936 | 184 | 7.5 | 18 | 93 | 235 | 11 |
| Median fold change | 1.3 | 1.6 | 2.3 | 1.6 | 1.4 | 1.5 | 2.1 |
| 2-tail MWU *P-value* | 1.9E-6 | 8.6E-7 | 3.3E-5 | 1.5E-3 | 1.3E-3 | 1.8E-3 | 1.2E-5 |

## DISCUSSION

Conventional serological assays detect antigen-specific polyclonal antibodies in serum using indirect immunoassays and report a ‘titer,’ defined as the reciprocal of the highest dilution of a patient sample that produces a positive signal. Measurements of polyclonal antibodies can be highly variable and depend not only on patient characteristics, including immunocompetence, time after pathogen exposure, and cross-reactivity from previous infections^49^, but also on calibration standards, interpretation of assay results, and reagent quality, including the stability, purity, glycosylation, and antigen conformations. In the absence of universal polyclonal antibody standards for individual pathogens and antigens, each diagnostic laboratory often uses its own set of ‘high, ‘medium,’ and ‘low’ titer samples, without inter-laboratory standardization. While monoclonal antibodies have been suggested as calibrators of serological testing^50^, they may not adequately recapitulate the equilibrium and binding kinetics of the complex mixture of isotypes and clones that differ in epitope specificity, concentration, affinity, binding kinetics, and competitive binding properties. Consistent with these limitations, substantially different levels of anti-RSVF IgG antibodies have been reported in adult sera, ranging from 4 µg/mL^12^ to 30 µg/mL^51^ to 300 µg/mL^13^. Such vast differences may highlight the need for next-generation serological assays.

We previously demonstrated numerous applications of bottom-up proteomics for identification and quantification of human proteins^52–64^, discovery of protein biomarkers^65–75^, and investigation of the functional roles of human proteins^76–80^. More recently, we have focused on developing novel MS approaches for the characterization of endogenous human polyclonal antibodies and the differential quantification of antibody isotypes and subclasses^2, 3, 81, 82^. In this study, we developed ProWiSe assays to enable in-depth characterization of endogenous polyclonal antibodies against human pathogens^83^ and to improve serological testing.

We suggested that next-generation serological assays could substantially benefit from the use of reference standards that are chemically defined, stable, enable absolute quantification, can be synthesized at scale, and could serve as universal reference standards for calibrating existing laboratory-specific standards. In this study, we presented potentially universal peptide standards (**Supplemental Table S4**) and the next-generation serological assays that may enable inter-laboratory standardization of serological testing. Direct quantification of antibody heavy-chain constant regions eliminates the need for secondary detection antibodies and mitigates several limitations of indirect immunoassays, including detection-antibody cross-reactivity, interference from antibody-associated proteins and complexes (c1q, CD5L), allotypic variations, and glycosylation heterogeneity. Furthermore, exhaustive denaturation, trypsin digestion, and a focus on a few defined tryptic peptides collapse the vast diversity of antibody clones and variants into a simple set of quantitative reference peptides.

Previous studies of acute viral infections have often shown a conventional Type 1 T helper (Th1) immune response, with Th1-type cytokines prompting class switching from IgM to IgG3, IgG1, and IgA1^84^. Conversely, chronic infections and repeated vaccinations were shown to induce a Type 2 T helper (Th2) or mixed Th1/Th2 response accompanied by class switching to IgG4, IgA2, and IgE^4, 5, 85^. Previous RSV studies have reported high titers of serum IgG1, IgG2, and IgG3, with substantially elevated anti-RSVF IgG1 and anti-RSVG IgG2 antibodies^11^. In this study, we hypothesized that the anti-RSV humoral response could be more complex than the acute SARS-CoV-2 humoral response^3^ and could be a useful model to evaluate ProWiSe assays and explore the diversity and dynamics of the anti-RSV immune response. ProWiSe assays revealed the involvement of nearly all human antibody isotypes and subclasses, except IgE, suggesting a mixed Th1/Th2 response. RSVG also elicited a diverse antibody response, but no RSVG-specific IgD or IgE were detected. It should be noted that IgE has not been detected by any targeted or exploratory untargeted assays in this study or in our previous studies^3^.

The untargeted ProWiSe assays enabled the identification of serum proteins co-enriched with human immunoglobulins, including CD5L, which has been reported covalently bound to serum IgM^23, 24^, the C1q complement complex subunits C1QA, C1QB, and C1QC, and complement C4B protein. Activation of C4B through cleavage of C4 protein by complement components and covalent binding of its reactive thioester groups to serine, threonine, and lysine residues on proteins on the surface was recently demonstrated^86^. The co-enrichment of the C1q complex and C4B suggested a potential role for anti-RSVF antibodies in complement-dependent cytotoxicity. Interestingly, soluble forms of Fc receptors^87^ were not identified. Collectively, ProWiSe workflows may serve as surrogate assays for investigating complement activation of human immunoglobulins.

The identification of secreted high-affinity IgD against RSVF in serum was the most unexpected finding from the ProWiSe assays. The functional role of IgD has been a mystery since its discovery in 1965^88^. Membrane-bound IgD, which is often co-expressed with IgM on the surface of naïve B cells, is an integral part of a B cell receptor complex and is widely used as a marker for B cell sorting^28, 29, 89, 90^. A specific Fc receptor for IgD has been suggested, but its identification remains controversial^28^. Total soluble IgD was reported in human adult serum at low levels (median 30 µg/mL)^39^ compared to other antibody isotypes and subclasses, and was in line with our measurements in maternal sera (median 87 µg/mL).

Antigen-specific IgD antibodies were previously identified against a small-molecule hapten^47^ and milk proteins^47^. A recent study sequenced membrane IgD of class-switched IgD+IgM-memory B cells, reconstructed the hypermutated IgD VDJ sequences as secreted IgG1 antibodies, and demonstrated that individual clones exhibited polyreactive binding to a panel of commensal and environmental antigens^91^. Secreted IgD was proposed as an amplifier of the Th2 immune response^47^ and an immunomodulator of self-reactive B cells^28, 92^. While some acute infections, such as rubella and measles, and chronic infections, such as hepatitis and malaria, were found to be associated with elevated total serum IgD^28^, to our knowledge, detection of a virus-specific high-affinity IgD circulating in serum has not been previously reported. A subset of regulatory B cells (CD19^+^CD5^+^CD10^-^CD3^-^) has previously been shown to secrete polyreactive RSVF-specific IgM and was proposed as a prognostic factor for RSV severity^10^. The presence of secreted high-affinity anti-RSVF IgD in serum warrants further investigation into its clinical significance. The identification of high-affinity polyclonal IgD recognizing prefusion and postfusion RSVF forms may facilitate the development of next-generation neutralizing antibodies^6, 93, 94^ and vaccines^35^.

Secreted IgD and polymerized surface IgD shed from B cells were previously found to be associated with enhanced humoral immune responses and increased production of antigen-specific mouse IgM, IgG1, and IgE antibodies^46, 47, 89, 95–100^. In earlier studies, treatment of mouse B cells with anti-IgD antibodies led to shedding of polymerized surface IgD, which subsequently induced production of IgD-binding factors in T cells and resulted in elevated production of mouse IgM and IgG1 antibodies^95–100^. Elevated production of antigen-specific IgM and IgG was observed in mice with IgD-producing plasmacytomas^46^. Stimulation of the humoral immune response by secreted antigen-specific IgD in mice was linked to basophil activation, IL-4 secretion, modulation of a Th2-type response, and increased production of IgG, IgM, and IgE antibodies^47, 89, 101^. The evolutionary conservation of IgD and its ability to engage basophils suggest that IgD may function as part of an ancestral immune-surveillance mechanism that enhances immune responses^48^.

Further development of ProWiSe assays will enable characterization of functional repertoires of antigen-specific polyclonal antibodies circulating in serum. Such diversity could previously be estimated primarily at the transcript level through single-cell RNA sequencing of B cell receptors^1^. The major limitation of that approach includes sequencing of memory B cells obtained from peripheral blood rather than antibody-secreting long-lived plasma cells of bone marrow^102, 103^. Other limitations included inability to predict concentration, affinity, and diversity of the functional antibody clones circulating in serum. Our ProWiSe assays provided a protein-level view of IGHV gene usage in the polyclonal antibody response to RSV, identifying 47 IGHV genes and suggesting that a limited subset of IGHV germline genes may dominate the serum polyclonal antibody response. Collectively, our findings highlight the complexity and diversity of the humoral immune response to RSV.

## CONCLUSIONS

In summary, ProWiSe assays provide a direct and comprehensive approach for the identification, absolute quantification, and characterization of serum polyclonal antibodies, including the detection of rare antibody isotypes, profiling the variable chain repertoires, and discovery of immunoglobulin-associated interactomes. ProWiSe assays can be adapted to investigate a diverse range of antigens and epitopes, biological fluids, and diseases, thus providing a framework for quantitative serology testing with potential applications in biomedical research and clinical diagnostics.

## ACKNOWLEDGMENT

We thank David Marchant, University of Alberta, for providing pre-stabilized prefusion RSVF (RSVF L305I DS-Cav1) protein.

## ABBREVIATIONS

iBAQ: intensity-based absolute quantification
LC-MS: liquid chromatography - mass spectrometry
LFQ: label-free quantification
MWU: Mann–Whitney U test
PBS: Phosphate Buffered Saline
RSV: respiratory syncytial virus
DDA: data-dependent acquisition
DIA: data-independent acquisition
SRM: selected reaction monitoring
post-iRSVF: post-fusion RSV glycoprotein F expressed in baculovirus-insect cells
post-hRSVF: post-fusion RSV glycoprotein F expressed in human HEK293 cells
pre-hRSVF: a stabilized prefusion RSV glycoprotein F expressed in human HEK293 cells
LOD: limit of detection

