## Supplemental Figures for "Proteome-Wide Serology Identifies Secreted IgD against RSV Fusion Glycoprotein as a Signature of Broadly Elevated Antibody Response"

**Supplemental Text**

EXPERIMENTAL METHODS

**Pre-stabilized RSVF.** Pre-stabilized RSVF (RSVF L305I DS-Cav1) was provided by the David Marchant Lab (University of Alberta). Briefly, L305I DS-Cav1 plasmids were generated by transforming high-efficiency NEB10-beta competent *E.coli* according to the NEB protocol*.* Plasmids were sent to Dr. Joanne Lemiuex’s lab at the University of Alberta, where the protein was expressed in Expi293F cells. Five days following transfection, cells were pelleted, and the protein expression was confirmed by western blot. The protein was purified on an AKTAstart (Cytiva Life Science, Cat # 29022094) using a HisPur Ni-NTA chromatography cartridge (Thermo Fisher Scientific, Cat # 90099) under isocratic elution conditions. The protein was then concentrated and buffer-exchanged using an Amicon Ultra-15, 10K MWCO (MilliporeSigma, Cat # UFC901024). Final concentration was determined using the Qubit Protein Assay kit (Invitrogen, Cat # Q33212), and protein was stored at -80°C until use.

**Supplemental Figures**

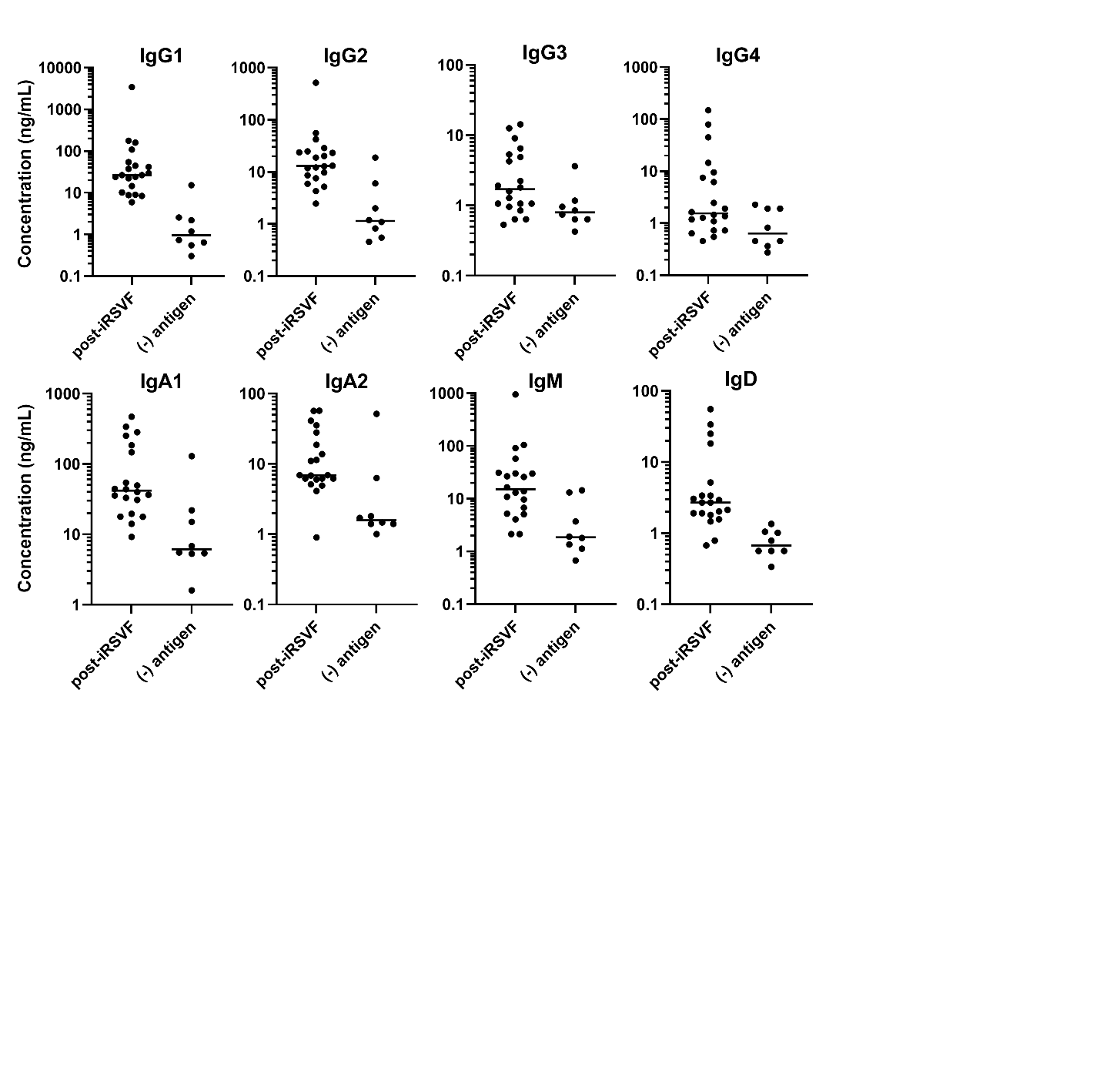

**Supplemental Figure S1.** **Quantification of endogenous polyclonal anti-post-iRSVF antibodies in male serum by rapid targeted LC-SRM.** Polyclonal antibodies were enriched from 20 healthy male serum using post-iRSVF antigen, with 10 PBS no-antigen controls include to assess non-specific binding across all isotypes and subclasses. Heavy isotope-labelled peptide standards with trypsin-cleavable tags (SpikeTides_TQL) enabled ‘absolute quantification’ based on the light-to-heavy ratios derived from MS/MS area of selected isotype-specific SRM fragment ions.

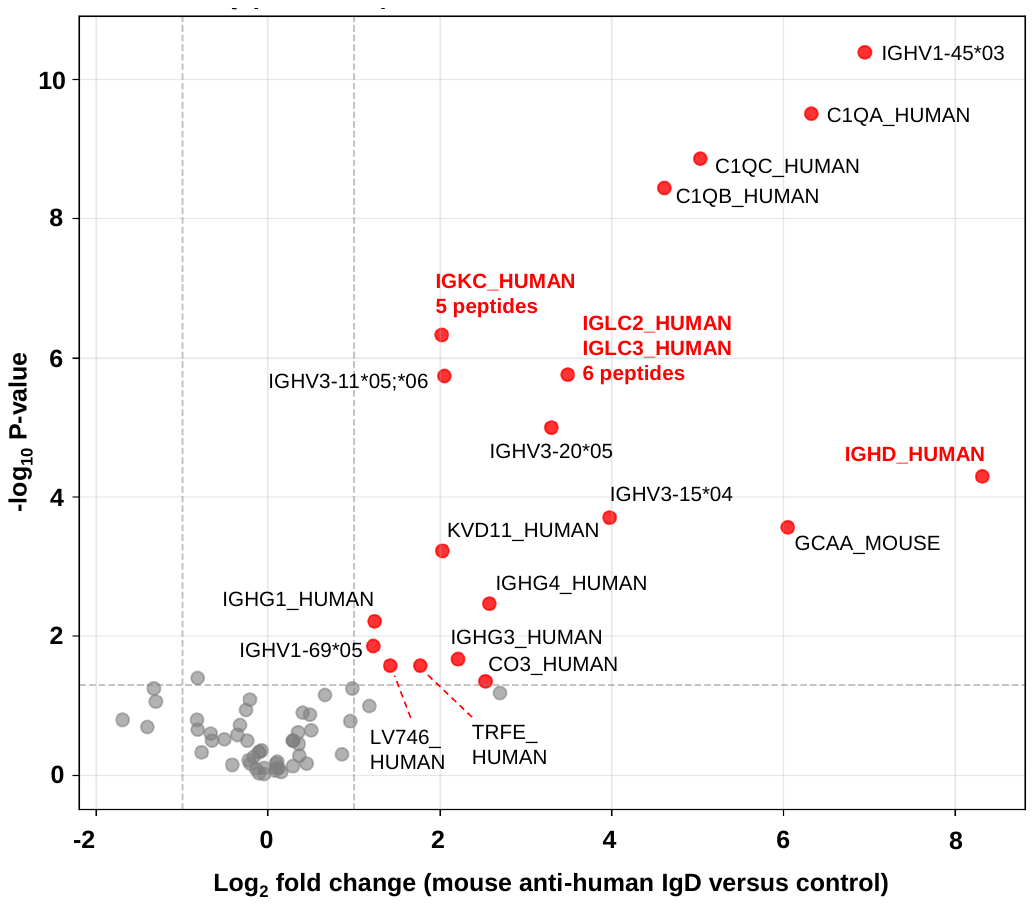

**Supplemental Figure S2. The usage of lambda and kappa light chains in secreted IgD.** Mouse anti-human IgD antibody was used to enrich IgD from serum. The vertical and horizontal dashed lines indicate the significance cutoffs (MWU P-value <0.05 and log2 fold change above +1 and below -1, respectively). IgD was immunoenriched 440-fold relative to negative control. The most abundant serum antibody isotype, IgG1 (typical serum IgG1/IgD ratio ~100) was enriched non-specifically by only 2.5-fold and therefore would not substantially contribute to κ/λ ratio. The κ/λ ratio for enriched IgD experiment was calculated as 0.59. For comparison, polyclonal antibodies enriched with post-iRSVF in our previous experiments, with IgG1 as the most abundant isotype, had κ/λ ratio of 1.78. These results indicate a preferential association of secreted IgD with λ light chains and are consistent with the previously observed “IgD paradox”.

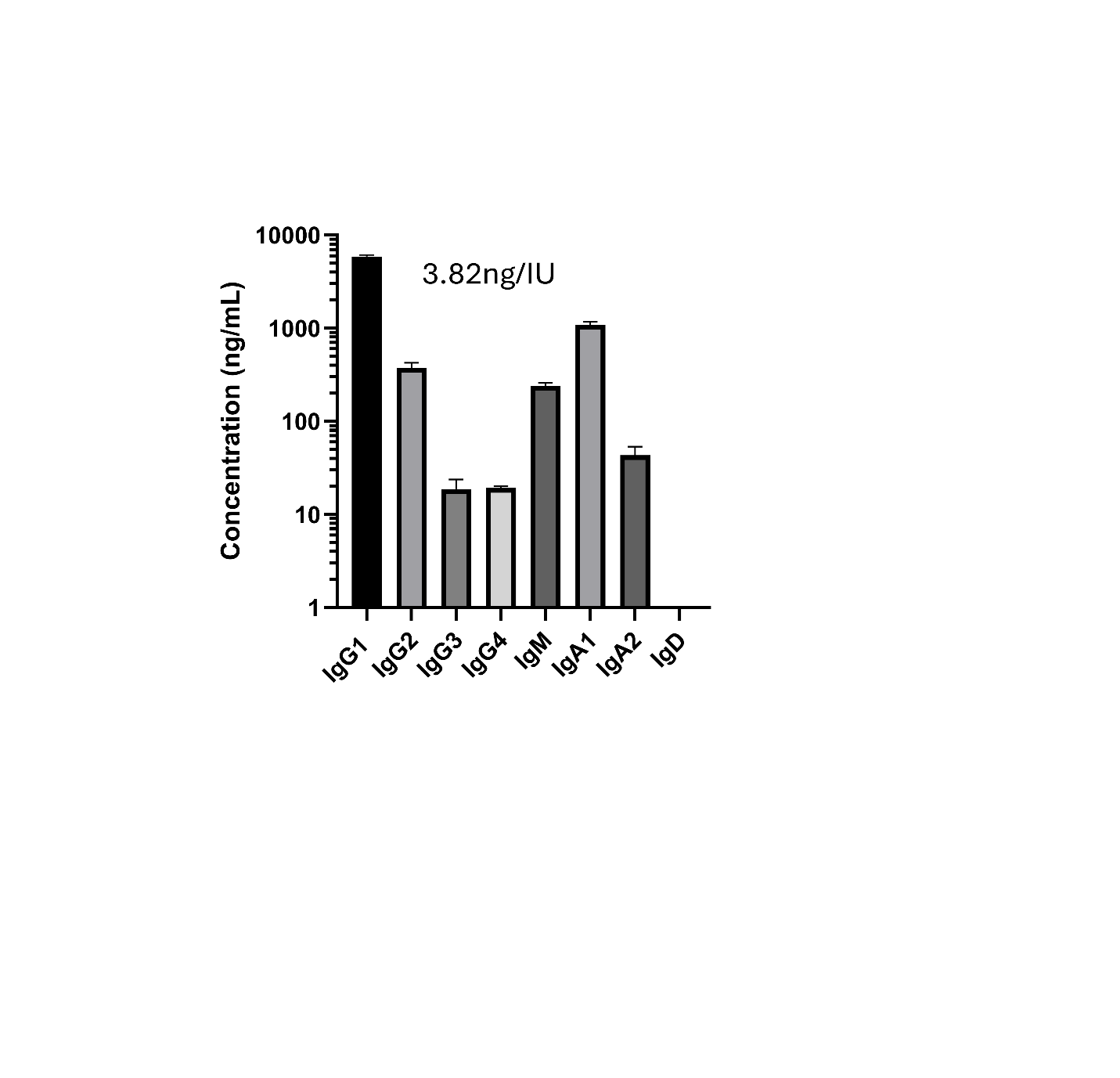

**Supplemental Figure S3. Concentrations of antibody isotypes and subclasses in the 1st WHO RSV Reference Serum.** Post-hRSVF was used to enrich RSVF-specific antibodies from WHO International Standard 16/284 for antiserum to Respiratory Syncytial Virus (2000 IU/mL). Absolute concentrations were measured with the targeted ProWiSe assay. Based on the assigned anti-RSV neutralizating antibody activity of the reference serum, 1 IU corresponded to 3.82 ng of combined IgG1, IgG2, IgA1, IgA2 and IgM in this sample. IU, International Unit of anti-RSV neutralising antibodies.
